# Gene expression plasticity of the mammalian brain circadian clock in response to photoperiod

**DOI:** 10.1101/2024.02.16.580759

**Authors:** Olivia H. Cox, Manuel A. Gianonni-Guzmán, Jean-Philippe Cartailler, Matthew A. Cottam, Douglas G. McMahon

## Abstract

Seasonal daylength, or circadian photoperiod, is a pervasive environmental signal that profoundly influences physiology and behavior. In mammals, the central circadian clock resides in the suprachiasmatic nuclei (SCN) of the hypothalamus where it receives retinal input and synchronizes, or entrains, organismal physiology and behavior to the prevailing light cycle. The process of entrainment induces sustained plasticity in the SCN, but the molecular mechanisms underlying SCN plasticity are incompletely understood. Entrainment to different photoperiods persistently alters the timing, waveform, period, and light resetting properties of the SCN clock and its driven rhythms. To elucidate novel molecular mechanisms of photoperiod plasticity, we performed RNAseq on whole SCN dissected from mice raised in Long (LD 16:8) and Short (LD 8:16) photoperiods. Fewer rhythmic genes were detected in Long photoperiod and in general the timing of gene expression rhythms was advanced 4-6 hours. However, a few genes showed significant delays, including *Gem*. There were significant changes in the expression clock-associated gene *Timeless* and in SCN genes related to light responses, neuropeptides, GABA, ion channels, and serotonin. Particularly striking were differences in the expression of the neuropeptide signaling genes *Prokr2* and *Cck*, as well as convergent regulation of the expression of three SCN light response genes, *Dusp4*, *Rasd1*, and *Gem*. Transcriptional modulation of *Dusp4* and *Rasd1,* and phase regulation of *Gem,* are compelling candidate molecular mechanisms for plasticity in the SCN light response through their modulation of the critical NMDAR-MAPK/ERK-CREB/CRE light signaling pathway in SCN neurons. Modulation of *Prokr2* and *Cck* may critically support SCN neural network reconfiguration during photoperiodic entrainment. Our findings identify the SCN light response and neuropeptide signaling gene sets as rich substrates for elucidating novel mechanisms of photoperiod plasticity.

## INTRODUCTION

The mammalian brain’s endogenous 24-hour timing mechanism, or circadian clock, is located within the suprachiasmatic nuclei (SCN) of the hypothalamus. SCN circadian clock neurons exhibit self-sustained circadian rhythms in gene expression which are driven by networks of clock genes organized in transcription/translation negative feedback loops (TTFLs, for review see Hastings et al., 2018). A key function of circadian clocks is to synchronize, or entrain, internal rhythms to local environmental time.

While much is known about the genes, neurons, and synapses that are critical for the generation of circadian rhythms, there are key gaps in knowledge regarding the molecular mechanisms of entrainment and the pacemaker plasticity that entrainment induces. Entrainment to different seasonal day lengths (photoperiods), or even a clock reset by a single light pulse, persistently alters the timing, waveform, period, and resetting properties of the SCN circadian clock and its driven rhythms (Buijink et al., 2016; Ciarleglio et al., 2010; Glickman et al., 2012; Inagaki et al., 2007; Pittendrigh et al., 1984; Pittendrigh and Daan, 1976a; Rohr et al., 2019; Tackenberg et al., 2020; VanderLeest et al., 2007, 2009). The long-term plasticity of circadian clocks in response to entraining light cycles is thought to be critical for stable alignment to changing seasonal photoperiods (Pittendrigh and Daan, 1976b, 1976c). Misalignment of the clock to the external environment may contribute to mood disorders, SCN changes in aging, and to discordance between SCN and peripheral tissue clocks (Baron and Reid, 2014). Here we describe the SCN transcriptomic response to different photoperiods to elucidate novel molecular mechanisms of SCN plasticity.

The molecular clock within SCN neurons is a nested multi-level oscillator. At its core is a two loop TTFL in which heterodimers of the transcription factors *Clock* and *Bmal1* positively drive the transcription of negative feedback *Period* circadian regulator genes (*Per1/2*) and *Cryptochrome* circadian regulator genes (*Cry1/2*) through E-box enhancer elements. PER and CRY proteins heterodimerize, accumulate, and enter the nucleus, inhibiting CLOCK/BMAL1-driven transcription, thus forming the core 24-hour feedback oscillator (For review see Hastings et al., 2018). In addition, the molecular circadian clockworks drives an extensive transcriptional network of clock-controlled genes in a tissue-specific and phase-specific manner, with nearly half the mouse genome thought to be under circadian control (Panda et al., 2002; Zhang et al., 2014).

The timing of the circadian clock molecular oscillator loop and its clock-controlled genes can be reset by light stimuli through synaptic input to SCN neurons from melanopsin-expressing retinal ganglion cells (Berson et al., 2002; Hannibal et al., 2002; Hattar et al., 2002; Provencio et al., 2002). The resetting transduction pathway within SCN neurons involves NMDA glutamate receptors and PAC1 PACAP receptors which depolarize the transmembrane potential and increase action potentials, raising intracellular Ca^++^ and cAMP levels, and ultimately stimulating MAPK/ERK and CREB/CRE transcriptional activation to induce rapid transcription of the *Period* genes (Obrietan et al., 1998, 1999). Importantly, secondary release of vasoactive intestinal polypeptide (VIP) from light-excited retinorecipient SCN neurons transmits light input broadly throughout the SCN neural network (Hamnett et al., 2019; Kuhlman et al., 2003), also through MAPK/ERK and CREB/CRE activation. SCN plasticity in response to light-induced clock resetting is thought to be critical to stable photoperiodic entrainment (Baron and Reid, 2014; Panda et al., 2002; Pittendrigh and Daan, 1976b; Rohr et al., 2019).

What then are the critical molecular mechanisms of the circadian clock photoperiod entrainment response? GABAAR signaling and its control by the chloride co-transporters, NKCC1 and KCC2, have been shown to be molecular elements in photoperiod plasticity in the SCN network (Farajnia et al., 2014; Myung et al., 2015; Rohr et al., 2019). However, these molecules alone are unlikely to fully account for the molecular bases of SCN plasticity, and the role of these transporters in the SCN has recently been called into question (Patton et al., 2023). To reveal additional candidate molecular mechanisms of SCN photoperiod plasticity, it is important to comprehensively define what changes occur in the circadian transcriptome of the SCN following photoperiod entrainment.

Here we have used RNAseq to characterize differential gene rhythmicity and differential gene expression in response to Short winter-like photoperiods vs. Long summer-like photoperiods. To best capture the impact of photoperiod, we used a strain of mice (C3Hf+/+, Baba et al., 2009) which possess intact melatonin signaling, and we used perinatal exposure to photoperiods, which we have previously shown to produce robust plasticity in the SCN and robust photoperiodic responses in downstream serotonergic and dopaminergic brain circuits (Ciarleglio et al., 2010; Giannoni-Guzmán et al., 2020; Green et al., 2015; Jameson et al., 2023; Siemann et al., 2019, 2020). Our approach reveals specific gene expression changes in SCN neuropeptide signaling and light response genes as candidate novel molecular mechanisms for SCN light response plasticity and SCN neural network reconfiguration in photoperiodic entrainment.

## METHODS

### Animals and Housing

The C3Hf^+/+^ mouse strain (a gift of Gianluca Tosini, Morehouse School of Medicine, Atlanta, GA) was used. C3Hf^+/+^ mice have intact melatonin signaling and do not carry the *rd* allele that causes retinal degeneration in the C3H/HeJ parent strain (Baba et al., 2009). Mice were paired and placed under Short (LD 8:16) or Long (LD 16:8) photoperiods (ca. 100 lux). Litters were born and maintained under these conditions. As rodent circadian rhythms develop in utero (Carmona-Alcocer et al., 2018; Davis and Gorski, 1988) during which maternal-fetal melatonin signaling can transmit photoperiodic information (Weaver and Reppert, 1986) and retinal melanopsin light transduction begins to function in the developing pups (Rao et al., 2013), the first litters born under the experimental photoperiods were not used in this experiment in order to avoid any potential after-effects of parental entrainment to previous photoperiod (standard institutional housing (LD 12:12)). Litters were raised with both the dam and the sire and were group housed, except for those placed in running-wheel cages for activity monitoring (see below). Mice were weaned at postnatal day 21 (P21) per standard practice and remained in the same photoperiod until postnatal day 50 (P50, late adolescence). All animal housing and procedures were approved by the Vanderbilt University Institutional Animal Care and Use Committee.

### Behavioral activity monitoring

A cohort of 10 mice (5 born and maintained under Short photoperiod and 5 under Long photoperiod) were singly housed and placed into cages with running wheels at P50. They were kept under the same LD cycle for a minimum of 9 days before being released into constant darkness (DD), during which they were allowed to freerun for at least 5 days. Wheel-running locomotor activity rhythms were recorded using ClockLab (version 6.1.06, ActiMetrics Software).

The phase angle of entrainment, Ψ (psi), was calculated as the timing of the activity onset in the first cycle in DD (from ClockLab) relative to the projected time of lights-off from the previous light dark cycle (projected Zeitgeber Time 12 [ZT12]). The daily duration of locomotor activity, α (alpha), was calculated separately for the LD and the DD conditions by first marking the onsets and offsets on each day in the actograms of each individual mouse using ClockLab.

For each individual mouse, the average onsets and offsets (h) for each day were calculated by averaging the points. Alpha was calculated as average offset – average onset. The free-running period of the locomotor rhythms, τ (tau), was determined using Lomb-Scargle periodogram analysis in Clocklab for the days in constant darkness (DD). Statistical analysis of Ψ, α, and τ values across photoperiods were performed with GraphPad PRISM.

### SCN dissection and RNA extraction

At postnatal day 50, the lighting regime was switched to constant darkness (DD) at the respective time of lights off (ZT 12) for each photoperiod. Mice spent a minimum of 36h in DD before sample collection began, such that gene expression in the samples likely represents the state of the endogenous clock rather than acute responses to the previously light cycle (Hughes et al., 2017). After this interval, SCN dissections took place at intervals of 4h over 2 circadian cycles (Hughes et al., 2017), until a minimum n of 4 was achieved per timepoint per cycle. Short and Long photoperiod samples were collected at 36, 40, 44, 48, 52, 56, 60, 64, 68, 72, 76, and 80 hours after lights off (Fig 1). A dim red headlamp was worn by the experimenter during the first steps of dissection. Mice were euthanized by cervical dislocation and the eyes removed to prevent potential light signaling to the SCN. The subsequent steps took place under light: the optic nerve was cut and the brain was removed such that it was situated ventral side up in a petri dish. The SCN was dissected from the brain under a microscope using Vannas-Tübingen Spring Scissors (15003-08, Fine Science Tools). Each SCN sample was put in a 1.5mL sterile DNase-RNase-free tube, and tubes were immediately placed in powered dry ice. RNA and DNA were extracted from each SCN sample using the AllPrep DNA/RNA Micro Kit (Qiagen) according to the instructions. (DNA was stored for later use).

**Figure 1.**
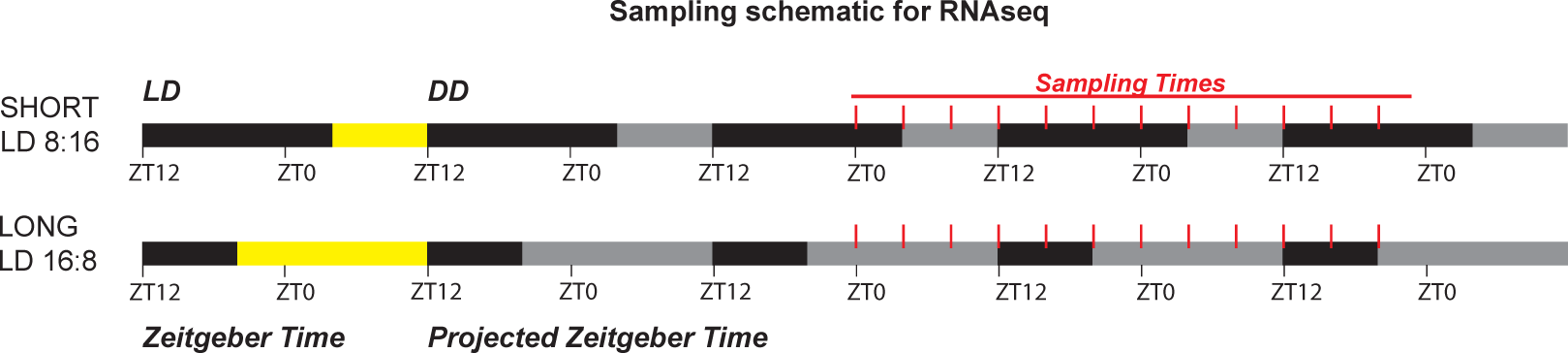
Sampling schematic for RNAseq. Last LD cycle before DD is shown at left in ZT, with yellow representing the light phase and where “lights off” is ZT12. At postnatal day 50, the lighting regime was switched to constant darkness (DD) at the time of lights off ZT12 as shown, (gray areas indicate the projected light phase), and mice spent a minimum of 36h in DD before sample collection, which occurred every 4h over 2 circadian cycles.

### RNA-sequencing

The quality of RNA samples was assessed using an Agilent 2100 Bioanalyzer or a PerkinElmer GX TOUCH 205 Nucleic Acid Analyzer. Only those samples with an RNA integrity number (RIN) of 7.9 or above were used for sequencing. Stranded cDNA libraries were constructed using 200 ng of total RNA per sample and the NEBNext® Ultra™ II RNA Library Prep (NEB, Cat: E7765S) per manufacturer’s instructions, with mRNA enriched via poly-A-selection using oligoDT beads. Paired-end sequencing produced approximately 54.6 million reads per sample (total of 11.1 billion reads), which were processed utilizing Trim Galore 0.6.7 (Krueger et al., 2023) and Cutadapt 1.18 (Martin, 2011) to remove adapter sequences and pairs that were either shorter than 20 bp or that had Phred scores less than 20. The Spliced Transcripts Alignment to a Reference (STAR v2.7.9a) application (Dobin et al., 2013) was used to perform sequence alignments to the mm39 (GRCm39) mouse genome reference and GENCODE comprehensive gene annotations (release M31). STAR’s two-pass mapping approach was used to increase the detection of reads mapping to novel junctions identified during the first mapping pass. Overall, 91% of the raw sequencing reads were uniquely mapped to genomic sites. RTA (version 2.4.11; Illumina) was used for base calling and demultiplexing. All data processing was performed at the Advanced Computing Center for Research and Education (ACCRE) at Vanderbilt University.

### Data Analysis

For downstream analysis, data from the 2 circadian cycles sampled were combined such that n ≥ 7 mice per timepoint. Differential analysis was performed by setting the Short photoperiod as the baseline in the differential gene expression pairwise comparisons. Since we used an mRNA library preparation to specifically capture protein coding genes, all analyses include and are normalized to protein-coding genes only (as determined by the gene_biotype obtained from BiomaRt).

### Differential expression analysis (DESeq2)

To investigate genes that were significantly up- or down-regulated in Long photoperiod as compared to Short, we used DESeq2 (Love et al., 2014). Of all 123 samples processed, 2 samples were identified as outliers and removed from downstream analysis. The design formula was ∼batch + photoperiod (12 sample preparations). Only genes that were present in at least 3 samples with normalized counts greater than or equal to 5 were retained. Gene annotations were retrieved from BiomaRt (Durinck et al., 2009) and mapped to Ensembl gene IDs. Genes without gene symbols were retained and labeled with their IDs.

### Differential rhythmic analysis (DiffCircaPipeline)

To explore differences in rhythmic biomarkers (e.g. differential rhythmicity, phase, amplitude, MESOR) between photoperiod groups the log2CPM expression data was analyzed using the DiffCircaPipeline v0.99.9 framework (Xue et al., 2023). Briefly, DCP uses a cosinor fit model (Cornelissen, 2014) to estimate from the time sample data amplitude (A), phase (Φ), Midline Estimating Statistic Of Rhythm (MESOR, M) and the noise level (σ) of gene expression rhythms. Unlike a cosine fit model, the cosinor fit model allows for the inclusion of additional parameters such as amplitude, acrophase (timing of the peak), and MESOR (mean level). This flexibility makes it more suitable for capturing variations in the shape and characteristics of the periodic data, unlike the simpler cosine fit model. First, DCP tests whether each gene is to be considered rhythmic in each condition (photoperiod) categorizing genes into four types of rhythmicity (TOR): arrhythmic in both groups (Arrhy), rhythmic in only group I (RhyI), rhythmic in only group II (RhyII), and rhythmic in both groups (RhyBoth) based on the cosinor fits. Then, DCP performs a differential rhythm fitness test for genes rhythmic in at least one group using the goodness of fit coefficient of determination R^2^. For genes determined rhythmic in both conditions, the framework also calculates differences in phase, amplitude and MESOR based on the cosinor fit. To account for the cyclical nature of phase (Φ) which DCP does not account for, 24hrs were subtracted from Δ peak phase values 12hrs or greater (Δ peak-24), while 24hrs were added to values less than or equal to −12hrs (Δ peak+24), effectively normalizing phase shifts to within a −12 to 12 range. We have reported in the main text findings by DCP q-value, but also report in the Supplemental Data DCP p-value for these parameters, which is comparable to CircaCompare output (Parsons et al., 2020). Our data was entered into the framework using median of ratios Log2 normalized data, which were corrected for batch effects using Limma (Ritchie et al., 2015), and then analyzed using the default settings.

### Gene Set Enrichment Analysis

To test for enrichment of functional gene groups in the DESeq2 differential expression data, DCP differential rhythmic amplitude data, and DCP differential MESOR data, we performed Gene Set Enrichment Analysis (GSEA, Subramanian et al., 2005a). For differential expression data (from DESeq2), log_2_fold change (< ± 0.10) and FDR adjusted p-value (<0.05) were used to create a ranked list. For differential rhythmic amplitude and differential MESOR (from DCP) p-value (<0.05) was used to create ranked lists. The C5 ontology gene sets collection of the Molecular Signatures Database (MSigDB) was queried (Liberzon et al., 2011, 2015).

### Phase Set Enrichment Analysis

To test for enrichment of functional gene groups with coordinated changes in entrained phase determine we used Phase Set Enrichment Analysis (PSEA) (Zhang et al., 2016). The input was the peak phases (Φ) of genes that were significantly rhythmic in both photoperiods as determined by DCP (S1, S3 Data). The default parameters for PSEA were used (Min genes per set = 10; Max sims/test =1000), and the C5 ontology gene sets collection (C5) of the Molecular Signatures Database (MSigDB) was queried (Liberzon et al., 2011; Subramanian et al., 2005b). PSEA evaluated each gene set to determine if the peak expression of these genes was coordinated in time using the Kuiper statistic (Kuiper.vs.Background (Zhang et al., 2016).

### Data Availability

Figures were generated using publicly available R-packages (The R foundation for Statistical Computing). All the code to generate the figures can be found (GitHub/Zenodo DOI available upon publication). Raw data has been deposited to NCBI’s Gene Expression Omnibus (#) and will be accessible upon publication. Analysis of differential expression and rhythmicity (phase, amplitude, MESOR) across photoperiods are available for the whole SCN transcriptome online.

## RESULTS

To confirm photoperiod entrainment and plasticity in the melatonin competent C3Hf+/+ strain, a cohort of mice were raised in Long (LD 16:8) or Short (LD 8:16) photoperiods and their locomotor activity rhythms were recorded in running wheel cages (see Methods). C3Hf+/+ mice released into constant darkness (DD) following entrainment to Long photoperiods exhibited significantly advanced activity onsets, significantly shortened activity duration, and trends toward shortened free-running period compared to mice entrained to Short photoperiods, similar to the original description of photoperiod plasticity effects in rodents (Pittendrigh and Daan, 1976a) (S1).

To identify candidate genes for SCN photoperiod-induced plasticity, we examined changes in the circadian transcriptome of the SCN from C3Hf+/+ mice raised in Long (LD 16:8) vs. Short (LD 8:16) photoperiods. Mice were paired and placed under Short (LD 8:16) or Long (LD 16:8) photoperiods and litters were born and maintained under these conditions. At postnatal day 50, the lighting regime was switched to constant darkness (DD) at the respective time of lights off (Zeitgeber Time 12 [ZT 12]) for each photoperiod. Mice spent a minimum of 36h in DD before sample collection began, and then SCN dissections took place at intervals of 4h over 2 circadian cycles (Fig. 1). We analyzed differential gene rhythmicity using DiffCircaPipeline (DCP) (Xue et al., 2023) and differential gene expression using DESeq2 (Love et al., 2014).

### Differential rhythmicity

Analysis of the whole SCN transcriptomic datasets with DiffCircaPipeline revealed that of the 16,549 transcripts detected by DCP, 2,488 genes, or 15% of the transcriptome, were detected as rhythmic in both photoperiods. 10,501 genes (63.5%) were considered arrhythmic, and 2,864 genes (17.3%) were found to be rhythmic in Short photoperiod only, while just 697 genes (4.21%) were found to be rhythmic in Long photoperiod only (Fig 2A). Fig 2B shows heatmaps of genes detected as rhythmic in both photoperiods by DCP. Analysis of the peak times of rhythmic genes as determined by DCP using cosinor fits showed two distinct groups in gene expression timing present in both photoperiods (Fig 2C), corresponding to the well-described “day” and “night” groupings in SCN gene expression (Panda et al., 2002). The timing of both gene expression peaks were advanced in Long photoperiod by ∼4-6h relative to Short photoperiod, which also is consistent with previous smaller scale studies of SCN gene expression in different photoperiods (Ciarleglio et al., 2010; Sumová et al., 2003; Tackenberg et al., 2020). Indeed, analysis of specific differential rhythmic parameters including peak phase, rhythmic amplitude, and the midline estimating statistic of rhythm (MESOR) (as illustrated in S2), demonstrated that the majority (73.5%) of genes that were rhythmic in both photoperiods had significantly different peak phases by q-value. The vast majority (1,825) of these genes were advanced in the Long as compared to the Short photoperiod, while only 4 genes were delayed: GTP binding protein overexpressed in skeletal muscle (*Gem*), HAUS augmin-like complex subunit 3 (*Haus3*), Leucine-rich repeats and IQ motif containing 3 (*Lrriq3*), and Ras association and DIL domains (*Radil*).

**Figure 2.**
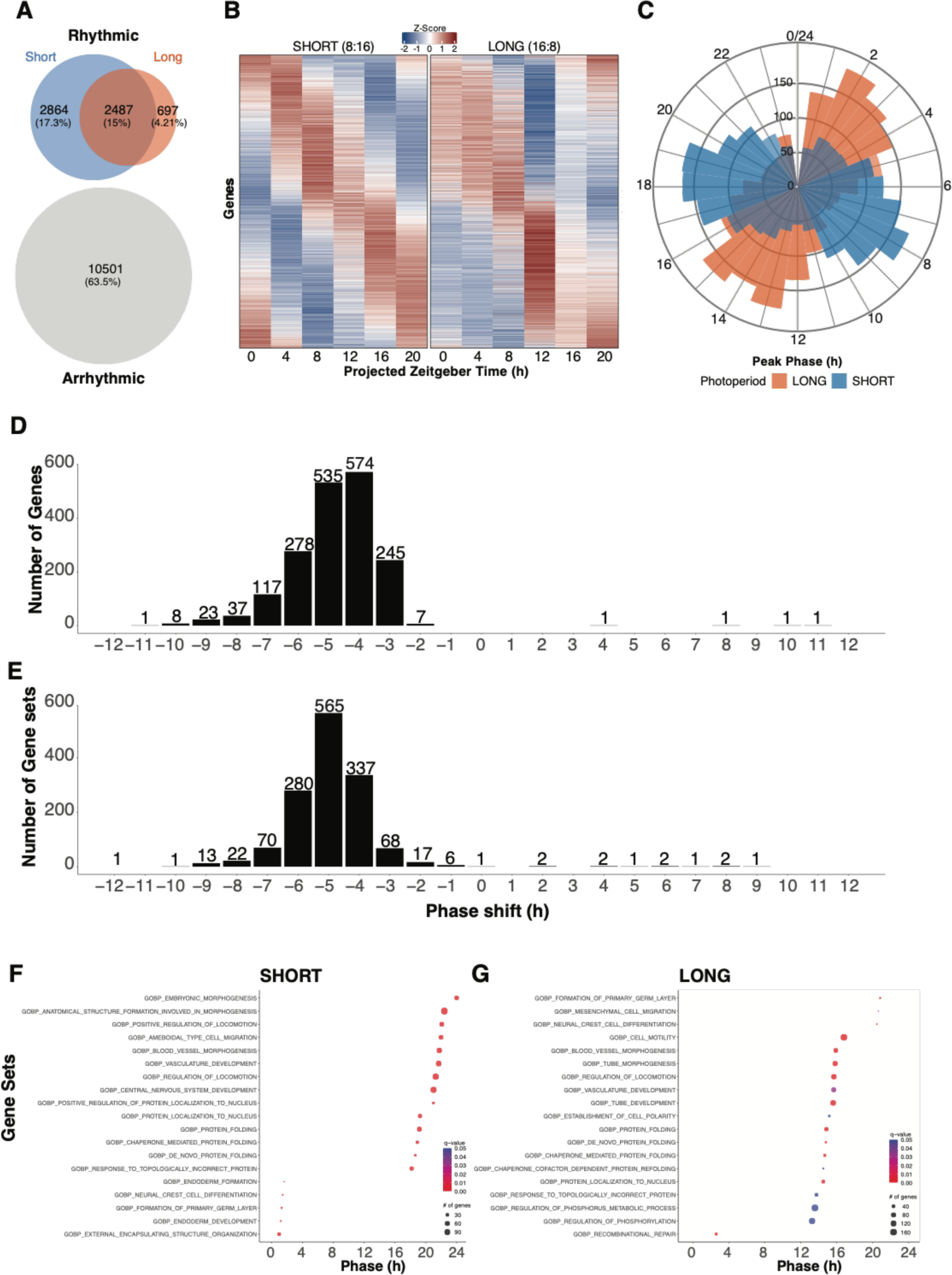
Transcriptome differential gene rhythmicity across photoperiods. **A)** Venn diagram showing the number of protein coding genes detected as rhythmic in Short, in Long, in both, and in neither photoperiod. **B)** Heatmap showing rhythmic gene expression in Short and Long photoperiods across time. **C)** Radar plot showing the peak phase for Short (blue) and Long (orange) photoperiods for genes determined to be rhythmic in both conditions. **D)** Histogram showing the distribution of genes according to their difference in peak phase (Long-Short). **E)** Frequency histogram delineating gene sets from the Gene Ontology database (PSEA) by their respective phase differences from Short to Long photoperiod. **F)** Column presenting the top 19 gene sets that exhibit significant phase coherence by the Kuiper statistic (q-value < 0.05) in short photoperiod. **G)** Column presenting the top 19 gene sets that exhibit significant phase coherence by the Kuiper statistic (q-value < 0.05) in Long photoperiod.

In contrast to peak phase, the rhythmic amplitude and the MESOR of genes were rarely found to be significantly different between photoperiods after DCP correction for false discovery rate (FDR) (q<0.05). However, it is worth noting here that previous analyses that employ cosinor fits to calculate the rhythmic parameters above, including the highly-cited “CircaCompare”, do not use FDR correction (Ding et al., 2021; Parsons et al., 2020). Therefore, we also investigated our rhythmic results at a threshold of p<0.05. 155 genes showed significant changes in rhythmic amplitude (1 increased, 154 decreased), while 447 genes showed significant changes in MESOR (209 increased, 238 decreased) in Long vs. Short photoperiods (S2, S1A Data). These DCP p-value significant changes are noted for individual rhythmic genes in the differential expression section below, and DCP output is also available parsed by the DESeq2 functional gene groups below (Circadian, Neuropeptide, Light Response, GABA, Ionic, Serotonin and Dopamine) in Supplemental Table S1B-G.

We also performed pathway analysis on the differential rhythms dataset using Phase Set Enrichment Analysis (PSEA, Zhang et al., 2016) and Gene Set Enrichment Analysis (GSEA, Subramanian et al., 2005b). PSEA of the genes detected as phase shifted in Long vs Short by DCP (q<0.05), revealed a broad peak of genes phase advanced centered at 4-6 hours (Fig 2D, S2 Data), in agreement with the absolute phases of gene expression plotted in Fig. 2C, as well as a similar broad peak for gene sets with shifted timing, again centered at 4-6 phase advanced (Fig 2E, S2 Data). Pathway enrichment for phase shifted genes in Short and Long photoperiods both included Regulation of Locomotion, which contains circadian-related genes, as well as other overlapping and non-overlapping gene sets (Fig 2 F,G; S2 Data). In addition, gene set enrichment analysis (GSEA) of genes that were significant for changes in rhythmic amplitude and MESOR showed pathway enrichment for Rhythmic Process and Positive Regulation of Gene Expression among others (Fig S3, S3 Data).

### Differential Expression

Analysis of differential gene expression using DESeq2 revealed salient differences in gene expression levels in the SCN transcriptomes of mice kept under Long versus Short photoperiods. Of the 16,856 transcripts detected by DESeq2, 1,518 were differentially expressed between the Short and Long photoperiods (p-adj < 0.05; Log2FC > [0.1]) (S4A Data). Of these, 972 were significantly downregulated and 546 were significantly upregulated in Long photoperiod. A volcano plot of the whole SCN data shows large magnitude changes in expression in known hypothalamic photoperiod response genes downstream of the SCN, such as *Lrat* in the retinoic acid signaling pathway (Shearer et al., 2010) and *Tshβ* (Korf and Møller, 2021), *Npvf* (Dardente and Simonneaux, 2022), *Dct* (Dardente and Lomet, 2018), *Dio2* (Korf and Møller, 2021), *Tafa3* (Korf and Møller, 2021), and *Slc16a2* (Ross et al., 2011) in the photoperiodic endocrine system (Fig 3), indicating that the photoperiod exposure indeed evoked a downstream photoperiodic response. Further, the direction of change agrees with multiple studies in mammals, with *Lrat*, *Npvf*, *Tshβ*, *Dct*, and *Dio2* all reported to be upregulated in Long photoperiod (Dardente and Lomet, 2018; Haugg et al., 2022; Korf and Møller, 2021; Shearer et al., 2010).

**Figure 3.**
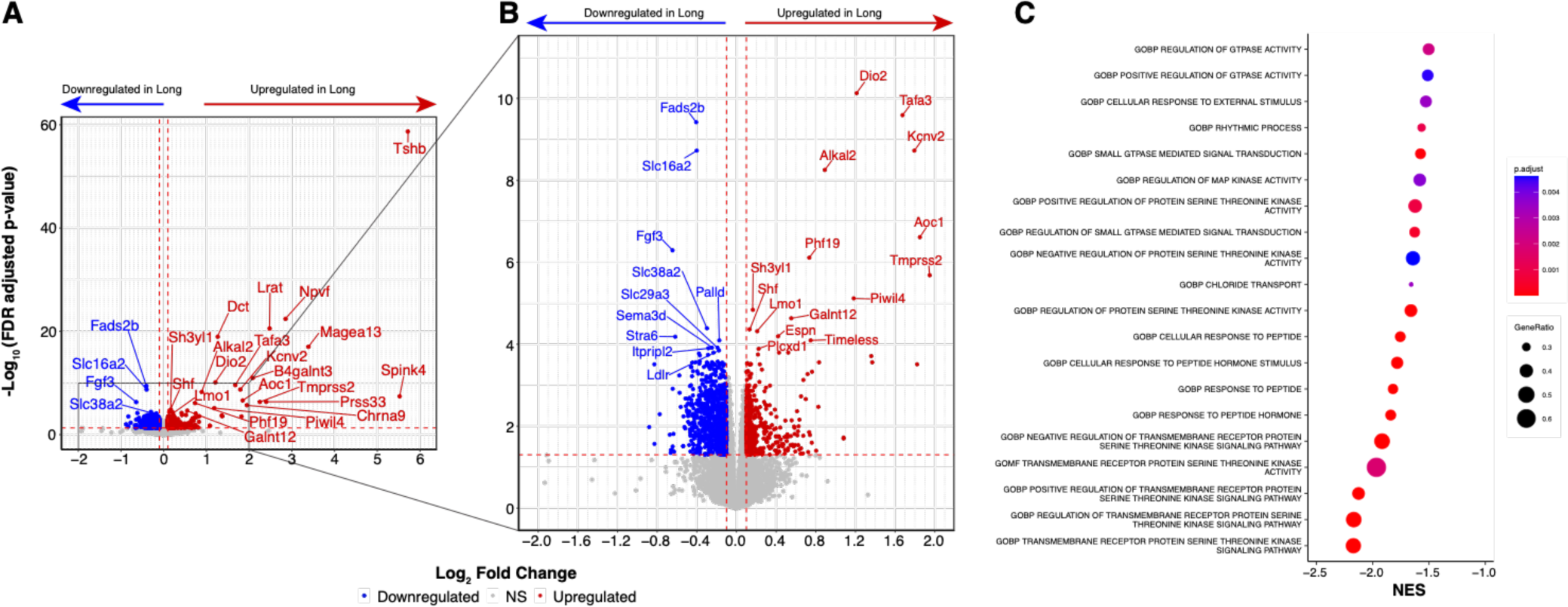
Transcriptome differential gene expression across photoperiods. **A)** Volcano plot showing changes in gene expression between Short and Long photoperiods. Directionality = from Short to Long (Long – Short). Downregulated genes are shown in blue and upregulated genes are shown in red. Thresholds for significance were set at log_2_FC ≥ ±0.1 (red dashed vertical lines) and p-adj < 0.05 (red dashed horizontal line). The top 25 genes meeting these criteria are labeled. **B)** Zoom view of volcano plot (grey box). Threshold for significance same as panel A. The top 25 differentially expressed genes in the plot are labeled. **C)** Pathway analysis showing relevant enriched gene sets, all of which were downregulated in Long photoperiod (as shown by negative normalized enrichment score (NES).

GSEA of differentially expressed genes revealed ca. 2400 enriched gene sets (padj<0.05) including 33 pathways related to locomotor rhythms, SCN light signaling pathways (GTPase, MAPK, S/T kinase), peptide hormone response, as well as GABA, chloride, potassium, and calcium transport (Fig 3C, S5 Data). To focus on genes robustly expressed in the SCN in our dataset, we used gene expression data from recent scRNA studies of SCN neurons for exploratory analysis (Morris et al., 2021; Wen et al., 2020). The gene sets from these studies have also been used to define SCN neurons as a class within the hypothalamus (Steuernagel et al., 2022). SCN neuron-expressed genes were enriched in SCN light response genes, neuropeptide signaling genes, and synaptic signaling genes, and we therefore further analyzed these SCN functional gene groups as described below (Figures 4-7, S5 and S6).

**Figure 4.**
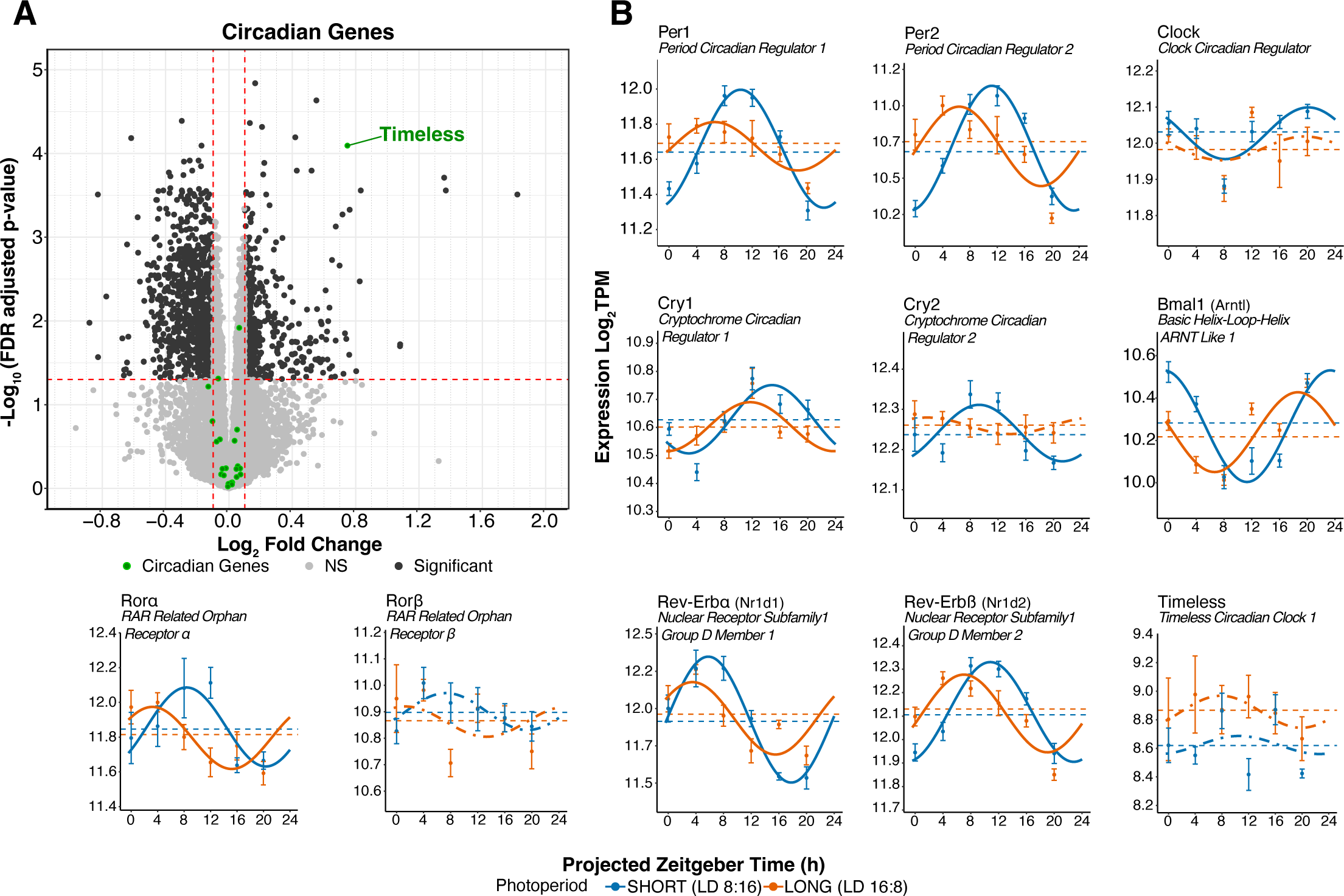
Circadian clock gene differential expression & rhythmicity in response to photoperiod. **A)** DESeq2 volcano plot of gene expression highlighting clock genes in green. Genes meeting the log_2_FC ≥ ±0.1 and p-adj < 0.05 are labeled. **B)** Cosinor plots illustrating the rhythmicity of selected clock genes. Solid lines depict significantly rhythmic cosinor fits, whereas dot-dashed traces indicate non-rhythmic fits. Dashed lines represent mean ±SEM for samples collected at each timepoint. Short (8:16) and Long (16:8) photoperiods are represented in blue and orange, respectively.

### Circadian clock genes

To determine whether photoperiod substantially impacted the core molecular clockworks, we analyzed differential expression and rhythmicity of 23 circadian clock genes and clock-associated genes detected by DESeq2. Overall, the expression and rhythmicity of these genes was consistent with expectations from previous studies (Panda et al., 2002; Zhang et al., 2014) (Fig 4). Surprisingly, we found no significant differences by DESeq2 in the expression level of core clock genes between Short and Long photoperiods. However, there was a significant increase in expression level of the circadian-associated gene *Timeless* in Long photoperiod (Fig 4A, S4B Data).

In terms of differential rhythmicity, the peak phases of the clock genes found to be rhythmic in both photoperiods (*Per1, Per2, Cry1, Arntl, Rorα, Nr1d1)*, were advanced 2-5 hours in Long photoperiod (DCP, q<0.05), similar to the overall pattern of phase advance of gene expression (Fig 4B), and the core clock gene *Per1* showed a significant loss of rhythmic power in Long photoperiod (DR fitness (ΔR^2^), DCP, q<0.05, S6 Data). Three additional clock genes, *Per2*, *Ciart*, and *Clock* showed trend level decreases in rhythmic power in Long photoperiod as well (DR fitness (ΔR^2^), DCP, p<0.05, S6 Data). In addition, *Per1, Per2, Arntl, Nr1d1*, and *Ciart* showed significant reductions in rhythmic amplitude by DCP p-value (<0.05), however none showed significant changes in MESOR (S1B Data). *Per1* and *Ciart* also showed decreased rhythmic amplitude in Long photoperiod at DCP q<0.10 (S1B Data). *Cry2* and *Clock* were detected by DCP as rhythmic only in Short photoperiod (S6 Data), whereas *Rorβ* and *Timeless* were not detected as rhythmic in either photoperiod.

### Neuropeptide signaling genes

Several neuropeptide signaling axes exist within the SCN neural network, including Arginine vasopressin (AVP), Cholecystokinin (CCK), Gastrin Releasing Peptide (GRP), Neuromedin S (NMS), Neuropeptide Y (NPY), Prokineticin 2 (PROK2), and Vasoactive Intestinal Polypeptide (VIP) (Hastings et al., 2018). A number of these peptides play important roles in SCN light responses, light resetting, and entrainment. In particular, VIP released from light-excited retinorecipient SCN neurons transmits light input broadly in the SCN neural network for light-induced resetting and neuronal synchrony (Hamnett et al., 2019; Kuhlman et al., 2003) and neurons expressing CCK are critical for responses to long photoperiods and may activate VIP neurons specifically under long photoperiod (Xie et al., 2023). We investigated 26 genes detected by DESeq2 that either make up the signaling axes themselves (peptides, receptors) or are involved in their modulation. Of these, we found that prokineticin receptor 2 (*Prokr2*), neuropeptide Y receptor Y1 (*Npy1r*), and neuromedin U receptor 2 (*Nmur2*) were significantly down-regulated in Long photoperiod, while cholecystokinin (*Cck*) and dual specificity phosphatase 4 (*Dusp4*), a modulator of VIP responses (Hamnett et al., 2019), were significantly up-regulated (Fig 5A; S2C Data). Rhythms in *Npy1r*, *Nmur2*, and *Dusp4* expression showed significant differences in peak phase between photoperiods (DCP, q<0.05). *Npy1r*, *Nmur2* also showed significant decreases in MESOR, while *Dusp4* showed a significant increase in MESOR (DCP p<0.05), however none of these genes showed significant changes in rhythmic amplitude by DCP p-value (Fig 5B, S1C Data).

**Figure 5.**
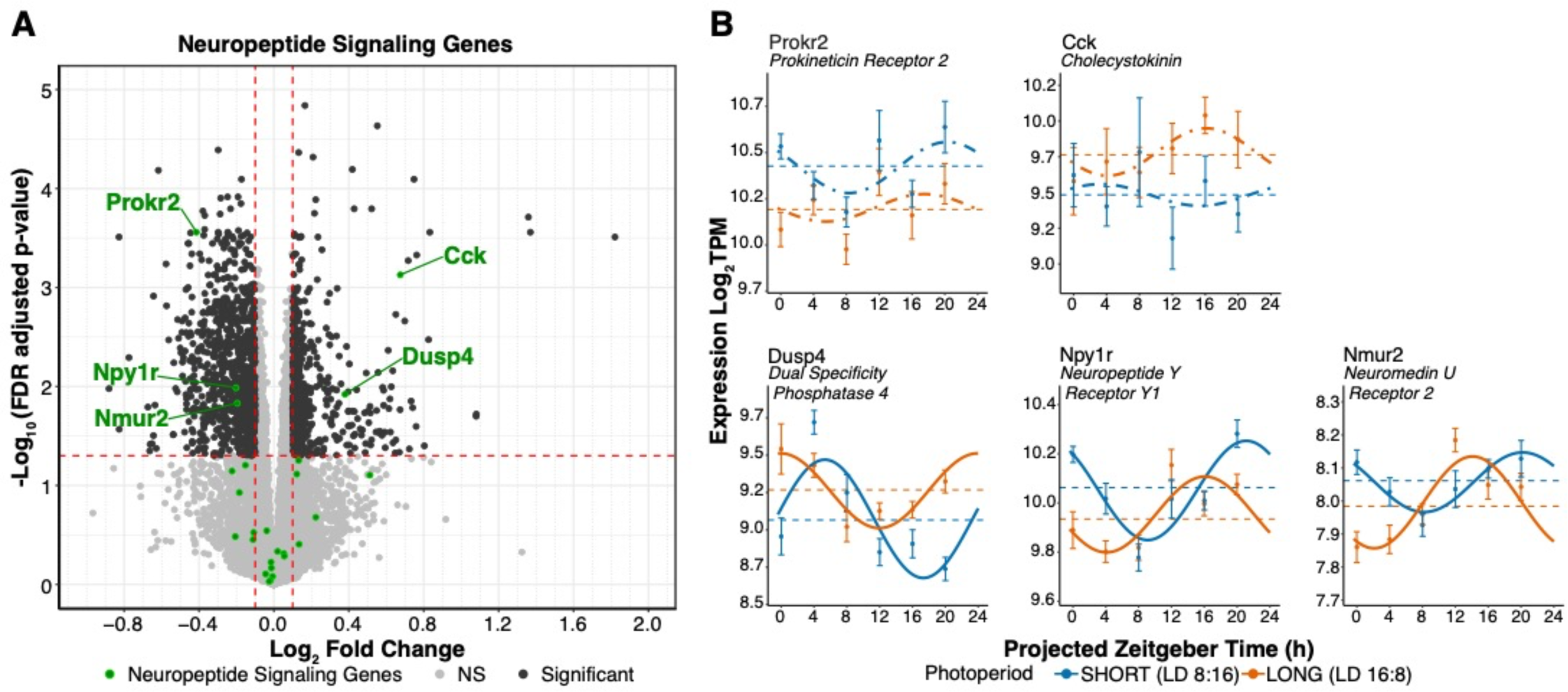
Neuropeptidergic signaling gene differential expression & rhythmicity in response to photoperiod. **A)** DESeq2 volcano plot of gene expression highlighting neuropeptide signaling genes in green. Genes meeting the log_2_FC ≥ ±0.1 and p-adj < 0.05 are labeled. **B)** Cosinor plots of differentially expressed genes. Solid lines depict significantly rhythmic cosinor fits, whereas dot-dashed traces indicate non-rhythmic fits. Dashed lines represent mean ±SEM for samples collected at each timepoint. Short (8:16) and Long (16:8) photoperiods are represented in blue and orange, respectively.

### Light responsive genes

Persistent alterations in SCN clock period are evoked by single clock-resetting light pulses as well as by entrainment to different photoperiods (Kim and McMahon, 2021; Pittendrigh and Daan, 1976a), and entrainment to different photoperiods alters the acute response of the SCN to light pulses (Glickman et al., 2012; Pittendrigh et al., 1984; VanderLeest et al., 2009), suggesting the possibility of mechanistic overlap between these responses to circadian light input. Therefore, we sought to examine whether the expression of acutely light responsive SCN genes might also be altered in our photoperiod treatments. A recent RNAseq study by Xu, et al. (Xu et al., 2021) examined which SCN genes responded acutely to light by exposing mice to pulses of light lasting 0.5-, 1-, 3-, and 6 hours. We compiled a global list of light responsive SCN genes in that study by combining the genes both up regulated and down regulated across all the pulse durations and removing those genes found in both groups. Of the 959 genes acutely regulated by light in the Xu et al., data (Xu et al., 2021), 99% of these genes were detected in our dataset, and 166 of these genes showed significant differential expression across photoperiods (S4D Data). Interestingly among the genes differentially expressed are *Dusp4* and *Rasd1* both of which are key modulators of the SCN light response through their actions on the critical NMDAR, MAPK/ERK and CREB/CRE light transduction pathway (Cheng et al., 2004; Hamnett et al., 2019) (Fig 6A). *Rasd1* was also detected as rhythmic in both photoperiods and showed the typical advanced peak phase in Long (DCP, q<0.05), as well as a decrease in MESOR and no change in rhythmic amplitude by DCP p-value (Fig 6B, p-value <0.05, S1D Data). DCP results for *Dusp4* are described in the neuropeptide signaling genes results. In addition, *Gem,* which also modulates SCN phase resetting to light through negative feedback onto the MAPK/ERK pathway (Matsuo et al., 2022), was one of the very few genes that showed a significant phase delay in Long compared to Short (Fig. 6B)(Carlson et al., 2016).

**Figure 6.**
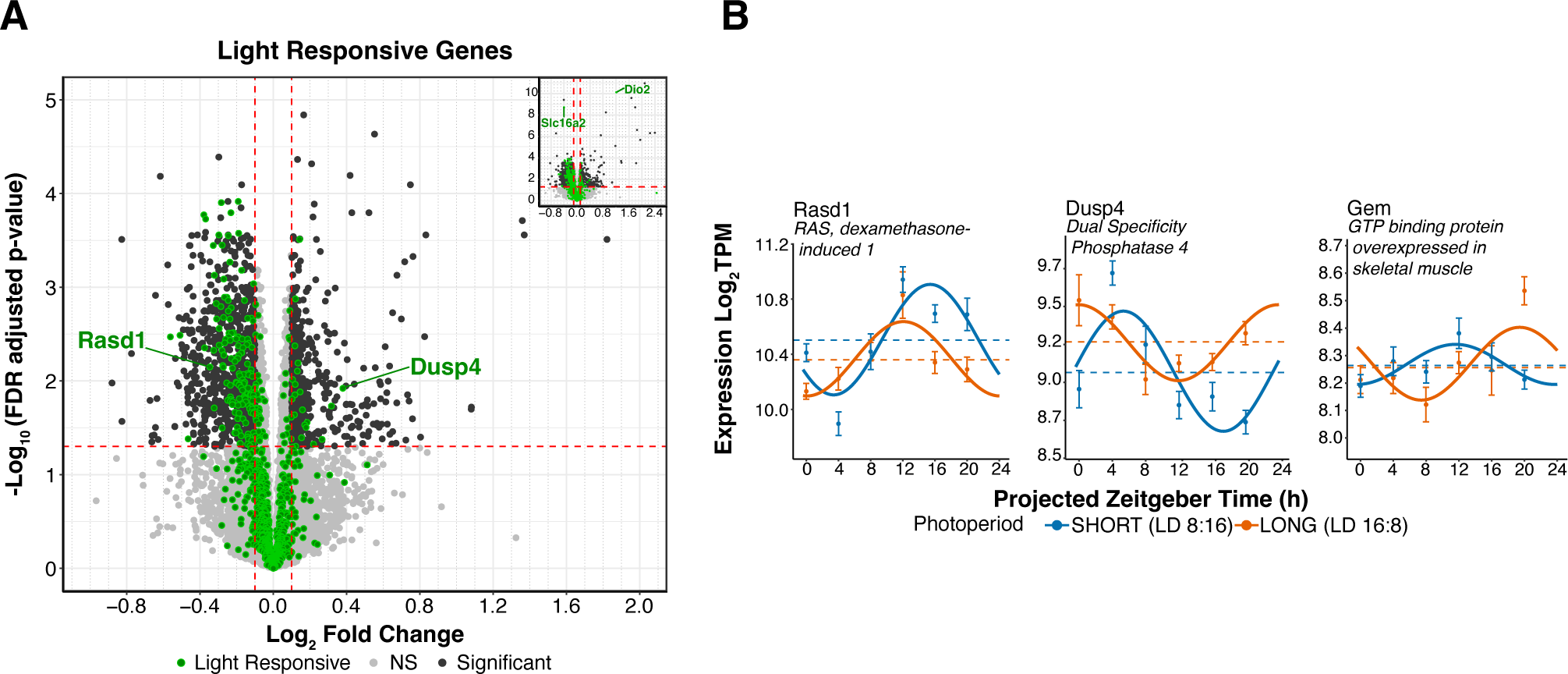
Light responsive gene differential expression & rhythmicity in response to photoperiod. **A)** DESeq2 volcano plot of gene expression highlighting light responsive SCN genes (Xu et al., 2021) in green at log2FC > ±0.1 and p-adj < 0.05. **B)** Cosinor plots of selected genes (described in A). Solid lines depict significantly rhythmic cosinor fits, whereas dot-dashed traces indicate non-rhythmic fits. Dashed lines represent mean ±SEM for samples collected at each timepoint. Short (8:16) and Long (16:8) photoperiods are represented in blue and orange, respectively.

### GABA signaling genes

Virtually all neurons in the SCN communicate via the neurotransmitter GABA, and photoperiodic regulation of inhibitory vs excitatory GABA transmission in the SCN is thought to be a key mechanism underlying photoperiodic plasticity (Farajnia et al., 2014; Myung et al., 2015; Rohr et al., 2019). Of 36 genes involved in SCN GABA signaling detected by DESeq2, 6 were found to be differentially expressed, all down regulated in Long photoperiod (Fig 7A, S4E Data). These included Na-K-2Cl cotransporter-1 (Nkcc1/Slc12a2), K-Cl cotransporter-1 (KCC1/Slc12a4), Betaine-GABA transporter (BGT1/Slc6a12/mGAT2), GABA transporter 2 (GAT2/Slc6a13/mGAT3), GABA_A_ receptor subunit rho 2 (Gabrr2), and GABA_A_ receptor subunit epsilon (Gabre) (Fig 7A). None of these genes were detected as rhythmic by DCP (Fig 7B).

**Figure 7.**
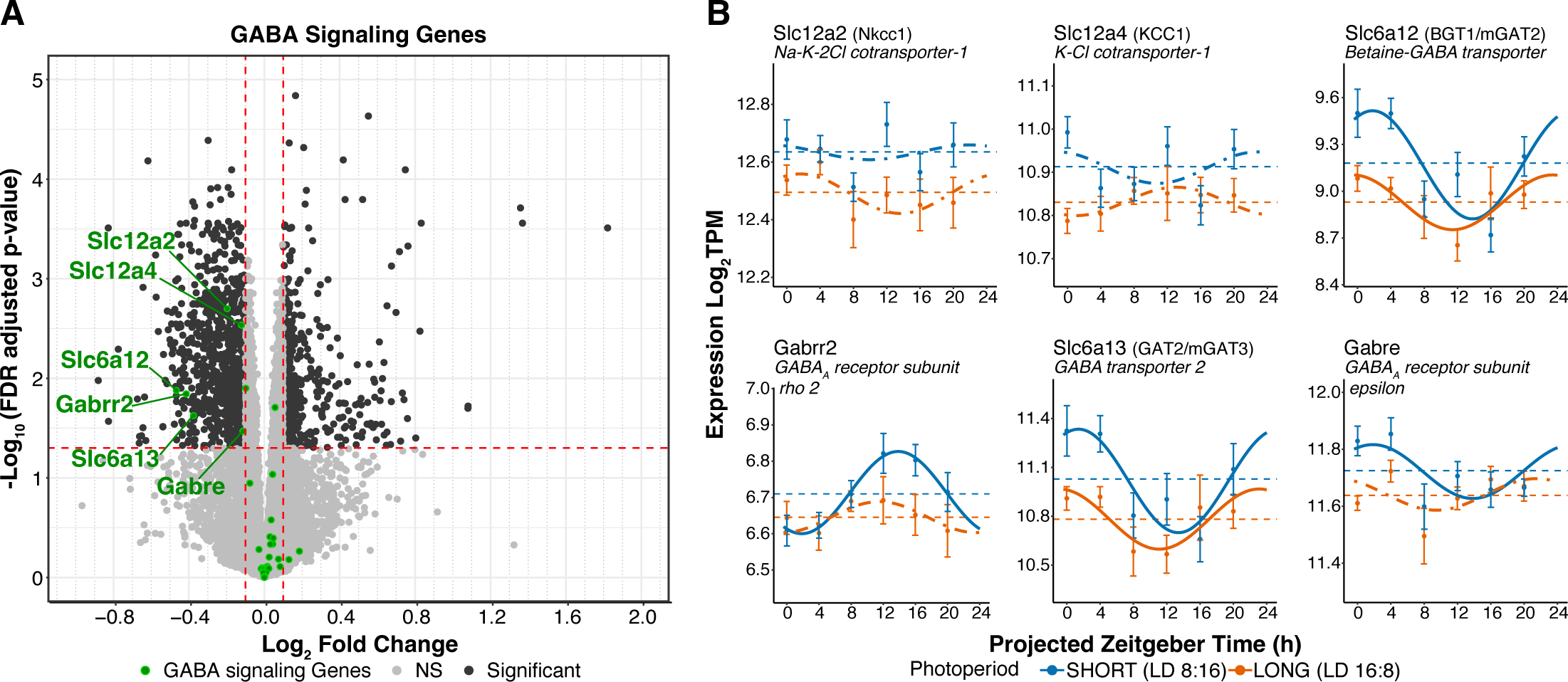
GABA signaling gene differential expression & rhythmicity in response to photoperiod. **A)** DESeq2 volcano plot of gene expression highlighting GABA signaling genes in green. Genes meeting the log_2_FC ±0.1 and p-adj < 0.05 are labeled. **B)** Cosinor plots of differentially expressed genes. Solid lines depict significantly rhythmic cosinor fits, whereas dot-dashed traces indicate non-rhythmic fits. Dashed lines represent mean ±SEM for samples collected at each timepoint. Short (8:16) and Long (16:8) photoperiods are represented in blue and orange, respectively.

### Ionic signaling genes

Previous work on SCN plasticity in response to entrainment to different light cycle periods has suggested the regulation of ionic and synaptic signaling genes as a potential mechanism for mediating that form of SCN plasticity (Azzi et al., 2017), therefore, we also examined calcium, potassium, sodium, and chloride voltage-gated channels genes, and related genes, in our dataset (166 genes total). Of these, 16 genes were differentially expressed between photoperiod conditions. 11 genes including *Kctd14, Kcnk5, Slc13a4, Kcnj13, Slc13a3, Kcna1, Ano1, Cacna2d4, Kcnk13, Kctd11, Kctd12* were down-regulated in Long photoperiod, while five genes including *Kcnv2, Scn4a, Clcn1, Sclt1, and Cacng7* were up-regulated in Long photoperiod (S4A; S4F Data). Of these *Kctd12* was detected as rhythmic in both photoperiod with phase advanced expression in Long (DCP, q<0.05), as well as decreased MESOR and no change in rhythmic amplitude by DCP p-value (S4B, p-value <0.05, S4B; S1F Data).

### Serotonin and dopamine signaling genes

Serotonin and dopamine inputs to the SCN modulate SCN light responses and entrainment (Grippo et al., 2017; Quintero and McMahon, 1999), and therefore we surveyed transcripts involved in serotonin and dopamine signaling in our dataset (24 genes total: 7 genes involved in dopamine signaling and 17 genes involved in serotonin signaling). The serotonin receptor 7 (Htr7) and the serotonin transporter (Slc6a4/Sert) were up-regulated, while the serotonin receptor 3B (Htr3b/5-HT3B) was down-regulated in Long photoperiod, while no dopamine signaling genes examined were found to have significant changes in expression (S5A; S4G Data). None of these genes were detected as rhythmic by DCP (S5B; S1G Data).

## DISCUSSION

Seasonal photoperiods entrain circadian rhythms and induce enduring plasticity in the SCN clock. To elucidate potential molecular mechanisms of photoperiod plasticity we performed RNAseq on whole SCN dissected from mice raised in Long (LD 16:8) and Short (LD 8:16) photoperiods. Overall, in terms of differential rhythmicity, fewer rhythmic genes were detected and there was an overall phase advance of gene expression rhythms of a few hours in Long photoperiod. Differential expression analysis showed no significant changes in the expression levels of the core clock genes, however, there were prominent differences across photoperiods in genes involved in SCN neuropeptide signaling and light response.

The advanced phase and decreased rhythmic power of core clock genes in Long photoperiod are consistent with previously demonstrated outcomes of entrainment to long photoperiods, including altered rhythmic waveforms and decreased rhythmic amplitude, as well as increased phase dispersal of single cell SCN rhythms (Buijink et al., 2016; Ciarleglio et al., 2010; Inagaki et al., 2007; Pittendrigh and Daan, 1976a; Sumová et al., 2003; Tackenberg et al., 2020; VanderLeest et al., 2007). In contrast, the clock-associated gene *Timeless* showed a significant increase in expression level in Long photoperiod. *Timeless* is a core clock gene in *Drosophila*, however its precise function in the mammalian SCN is unclear. Knockdown of *Timeless* abrogates SCN electrophysiological rhythms, while homozygous germ-line knockout is embryonic lethal, and heterozygotes show normal circadian rhythms (Barnes et al., 2003; Reppert et al., 2000). A human mutation in *Timeless* is associated with altered sleep phase and modifies clock period and stability of PER/CRY complexes in mice (Kurien et al., 2019). Thus, the photoperiod driven changes in *Timeless* expression we describe here have the potential to affect SCN rhythmic properties.

While photoperiod-induced changes in the expression of clock genes were relatively few, there were abundant changes in expression levels of SCN neural signaling genes related to neuropeptides, GABA, ion channels, serotonin, and dopamine. In all 29 genes from these categories were found to exhibit photoperiod-dependent changes in expression level using DESeq2. Previous results have established regulation of inhibitory vs. excitatory GABA signaling in the SCN through the expression of *Nkcc1* and *Kcc2* chloride transporter genes as a likely mechanism of photoperiodic plasticity (Farajnia et al., 2014; Myung et al., 2015; Rohr et al., 2019). Indeed, we found expression changes in six GABA signaling genes - Na-K-2Cl cotransporter-1 (*Nkcc1/Slc12a2*), K-Cl cotransporter-1 (*KCC1/Slc12a4*), GABA transporter 2 (*GAT2/Slc6a13/mGAT3*), GABA transporter BGT1 (*BGT1/Slc6a12/mGAT2*), GABA_A_ receptor subunit rho 2 (*Gabrr2*), and GABA_A_ receptor subunit epsilon (*Gabre*), all of which were reduced in expression in Long photoperiod. The expression of *Nkcc1* compared to *Kcc2* was previously found to increase in the dorsal SCN in Long days (Myung et al., 2015), however *Nkcc1* expression was reduced overall in our data, and *Kcc2* was not significantly changed. These differences may be due to the fact that photoperiod regulation of *Nkcc1/Kcc2* in the SCN is highly region-specific (Myung et al., 2015; Rohr et al., 2019), and our data is taken from whole SCN, perhaps obscuring regional changes. In addition, while we found expression changes in the GABA transporters *GAT2* (*Slc6a13/mGAT3*) and *BGT1* (*Slc6a12/mGAT2*), we did not detect significant changes in the principal SCN-expressed GABA transporters of *GAT1 (Slc6a1, mGAT1)*, which is expressed in neurons and astrocytes, and *GAT3* (*Slc6a11, mGAT4),* which is expressed in astrocytes and is critical for regulation of extracellular GABA levels and rhythms (Patton et al., 2023). Thus, the changes in expression of GABA signaling genes we observed present the possibility of contributing to SCN plasticity, but in themselves they do not establish a very coherent picture.

A recent report by (Porcu et al., 2022), described changes in the number of SCN VIP expressing and NMS expressing neurons in response to photoperiod. We did not observe differential expression of these peptide genes, but we did observe a significant change in the expression of *Nmur2*, the principal receptor for NMS in the SCN. There are substantial differences in both the mouse strains and photoperiod stimuli used in these studies that likely account for these apparent differences in results. Here we have used melatonin competent C3Hf+/+ mice and long-term entrainment to photoperiods including development (E0-P50), while Porcu et al., used melatonin synthesis deficient C57BL6 mice and 15 days of photoperiod entrainment in adult mice 9-16 weeks of age (Porcu et al., 2022).

A category of SCN genes that showed a particularly robust response to photoperiod were SCN light responsive genes – genes that change expression rapidly (within 6 hours) in response to a phase resetting light pulse (Xu et al., 2021). Remarkably, 166 acutely light responsive genes were found to exhibit significant changes in expression after weeks-long entrainment to Long vs Short photoperiods. It is important to note that these are sustained changes in expression in our experiments, rather than acute induction by light, because photoperiod entrained mice were maintained in constant darkness for at least 36 hours prior to sampling. Intriguingly, we found significant differential expression or rhythmicity across photoperiods of three acutely light responsive genes that are all modulators of the critical NMDAR, MAPK/ERK and CREB/CRE pathway for light response transduction in SCN neurons, *Dusp4*, *Rasd1*, and *Gem*. Thus, this light responsive gene set appears to be a particularly rich substrate for elucidating novel mechanisms of photoperiod plasticity in the future.

What are the specific changes in SCN function following photoperiodic entrainment that beg molecular explanation? Entrainment of rodents to long photoperiods (>12hr light/day) has been shown to have four distinct and enduring after-effects - to advance the phase of behavioral, electrophysiological, and molecular rhythms (Pittendrigh and Daan, 1976a; Tackenberg et al., 2020; VanderLeest et al., 2009). To alter the waveform of behavioral, electrophysiological, and molecular rhythms by compressing the duration of activity (α) and correspondingly expanding the duration of electrophysiological and *Period* gene expression (Pittendrigh and Daan, 1976a; Tackenberg et al., 2020; VanderLeest et al., 2009). To shorten the free-running period of behavioral, and molecular rhythms (Pittendrigh and Daan, 1976a; Tackenberg et al., 2020). And to decrease the responsiveness of the SCN to light input and its principal synaptic mediator NMDA glutamate receptors (Glickman et al., 2012; Pittendrigh et al., 1984; VanderLeest et al., 2009).

How might the changes in gene expression we have observed underlie these specific functional aspects of SCN photoperiod plasticity? Focusing on genes with robust changes in expression or rhythmic parameters (DESeq2 padj<0.05, Log2FC > [0.3], (Xu et al., 2021); DCP q<0.05) and for which there is evidence in the literature of SCN expression and function, there are several genes in our dataset that are of potential mechanistic importance. These include *Dusp4*, *Rasd1,* and *Gem* which are each key modulators of the SCN light response (Cheng et al., 2004; Hamnett et al., 2019; Matsuo et al., 2022). In addition, there were changes in the expression of *Prokr2* the widely expressed receptor for the critical PROK2 signaling network hub in the SCN (Morris et al., 2021), and in the expression of (*Cck),* which is expressed in neurons critical for light-induced phase advances and entrainment to Long photoperiods (Xie et al., 2023). Transcriptional modulation of *Prokr2* and *Cck* may critically support SCN neural network reconfiguration in photoperiodic entrainment (Buijink et al., 2016; VanderLeest et al., 2007) that may underlie changes in phase angle of entrainment, activity duration, or period.

Particularly intriguing is the convergent differential expression of *Dusp4*, and *Rasd1,* as well as differential rhythmic phase of *Gem* to potentially impact the SCN light response mediated by NMDAR, MAPK/ERK and CREB/CRE induction of light responsive genes. *Dusp4* is a phosphatase that targets phospho-ERK and is a negative modulator of the MAPK/ERK pathway and the SCN phase shifting response to light (Hamnett et al., 2019). Here we found that *Dusp4* expression is increased in Long photoperiod, which would be predicted to decrease ERK phosophorylation and its stimulation of CREB/CRE to induce the *Period* genes. *Rasd1* is a GTPase positive mediator of NMDAR coupling to MAPK/ERK stimulation in the SCN and is a positive mediator of the SCN phase shifting response to light (Cheng et al., 2004). Here we found that *Rasd1* expression is decreased in Long photoperiod, which also would be predicted to decrease ERK activation and stimulation of CREB/CRE. In addition, *Gem*, a small G-protein negative modulator of voltage-dependent Ca++ channels, mediates negative feedback to MAPK/ERK transduction of light responses within SCN neurons through rapid light induction and then inhibition of Ca++ channels to limit MAPK/ERK activation. *Gem* is robustly induced by light at phases when its rhythmic expression in low but is weakly induced when its expression is elevated near its peak (Matsuo et al., 2022). Based on this temporal pattern, the alignment of *Gem* peak phase we observe in Short photoperiod (Fig. 6B) would be predicted to decrease *Gem* light induction in the early night and thus enhance the amplitude of response to phase delaying light pulses, compared to *Gem’s* more delayed peak phase in Long.

Therefore, *Dusp4*, *Rasd1,* and *Gem* are all coordinately regulated in Long photoperiod in a manner expected to decrease MAPK/ERK activation, CREB/CRE driven *Period* gene induction, and the SCN phase-shifting light response compared to Short photoperiod. Indeed, reductions in the SCN response to NMDAR stimulation, in ERK phosphorylation, in *Per1* induction by CREB/CRE, and in behavioral phase resetting to acute light pulses have all been demonstrated in rodents following entrainment to long photoperiods (Glickman et al., 2012; Pittendrigh et al., 1984; VanderLeest et al., 2009). Transcriptional modulation of *Dusp4* and *Rasd1,* and phase modulation of *Gem* rhythms are therefore a compelling candidate molecular mechanisms for SCN light response plasticity. Interestingly, light sensitivity of the human circadian system, measured using acute melatonin suppression by light, is also decreased following exposure to long photoperiods vs short (Higuchi et al., 2007; Owen and Arendt, 1992), suggesting that similar mechanisms may impact the human circadian system. Notably, *Dusp4*, *Rasd1*, and *Gem* have demonstrated roles in hippocampal learning and memory and in cortical neuron plasticity (Abdul Rahman et al., 2016; Carlson et al., 2016; Takahashi et al., 2021) indicating potential mechanistic overlap between circadian clock plasticity and core mechanisms of neural plasticity in the mammalian brain (Buijink et al., 2016; VanderLeest et al., 2007).

In summary, using RNAseq on whole SCN dissected from mice raised in Long (LD 16:8) and Short (LD 8:16) photoperiods we found salient differences across photoperiods in the expression of genes involved in SCN light responsive genes and neural signaling genes. We describe a remarkable convergent regulation of three SCN light response and MAPK/ERK modulating genes - *Rasd1*, *Dusp4* and *Gem* - which potentially underlie plasticity of the SCN light response, and in addition, our dataset reveals several potentially novel SCN photoperiod plasticity genes including *Prokr2*, *Cck* neuropeptide signaling genes and the clock-associated gene *Timeless*. Future experiments can test the potential novel mechanistic roles of specific genes from this dataset in SCN photoperiod entrainment and plasticity using ex vivo optogenetic entrainment (Kim and McMahon, 2021) combined with SCN specific gene manipulation.

## ACKNOWLEDGEMENTS

Supported by NIH R01 GM117650 to DGM.

