## Supplementary Figures for "Gene expression plasticity of the mammalian brain circadian clock in response to photoperiod"

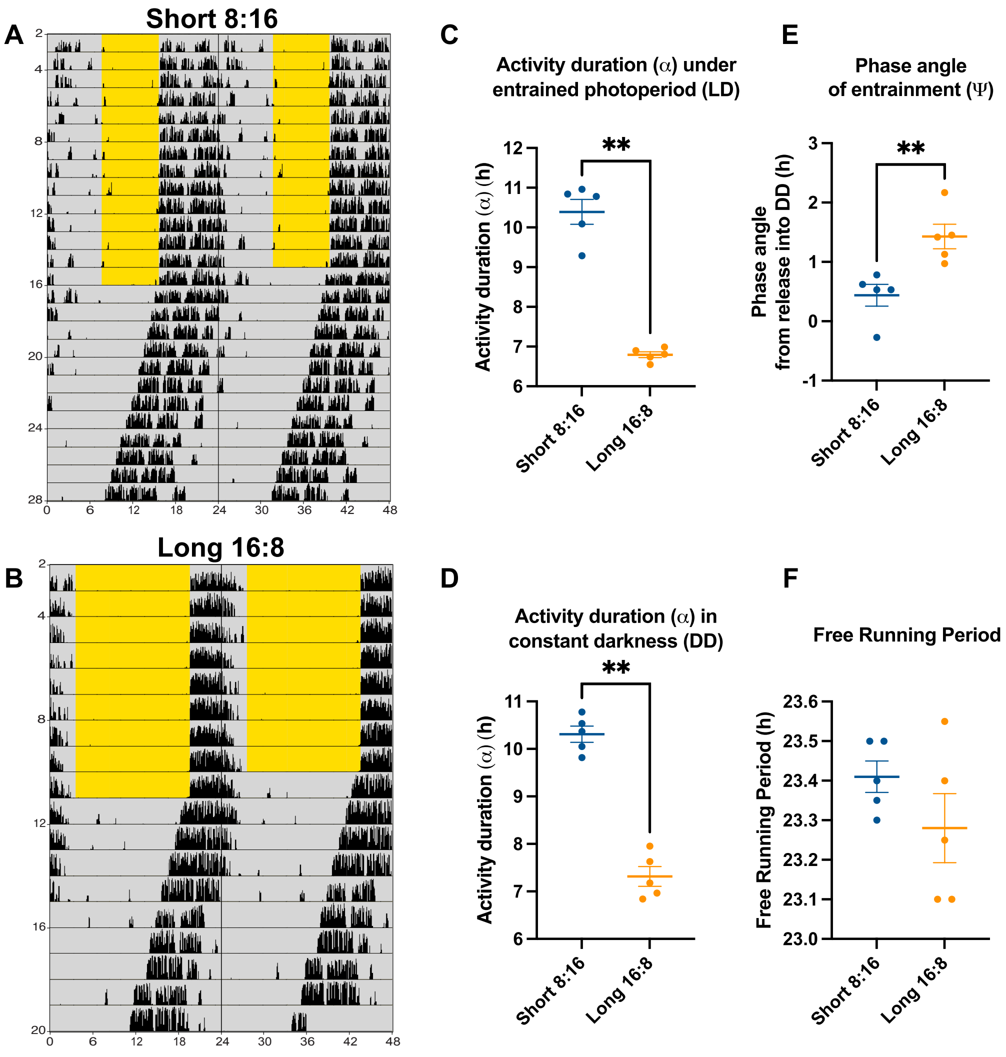


**Supplementary Figure 1. Effects of photoperiodic entrainment on the locomotor activity of C3Hf +/+ mice.** Double-plotted actograms for mice raised in **A)** Short 8:16 and

**B)** Long 16:8 photoperiods. After a minimum of nine days in a wheel running cage under light/dark cycle (LD) mice were released into constant darkness (DD). Activity duration (α) in **C)** LD (, **D)** DD, **E)** phase angle of entrainment (Ψ), and **F)** free running period were quantified for each individual (n=5 per group). Non-parametric Mann Whitney tests were performed to examine differences between Short and Long photoperiods in these parameters ($p < 0.01$** **C, D** and **E**).

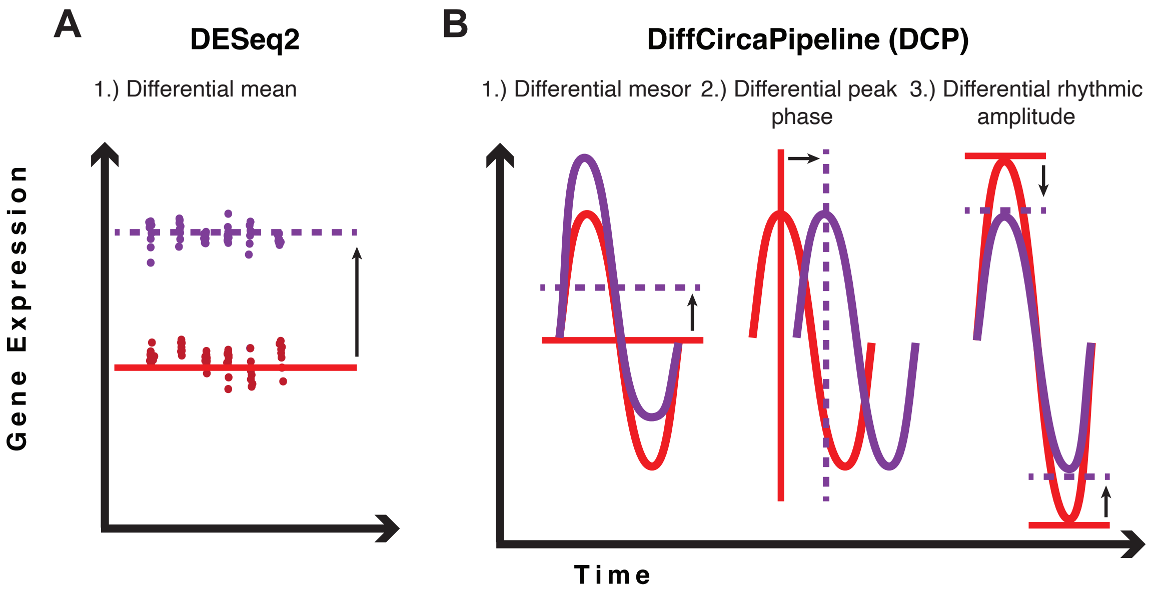


**Supplementary Figure 2. Types of gene expression changes observed in rhythmic genes. A)** By using DESeq2 we tested for overall differences in mean expression between each photoperiod by collapsing all the timepoints together **B)** In DiffCircaPipeline (DCP) we examined the following gene expression changes **1)** Differential expression represents an overall change in the mean expression (timepoints collapsed) between conditions. **2)** Differential peak phase represents a change in the peak expression time of a gene, as shown by a shift along the x-axis. **3)** Differential rhythmic amplitude represents a change in mean peak-to-trough amplitude.

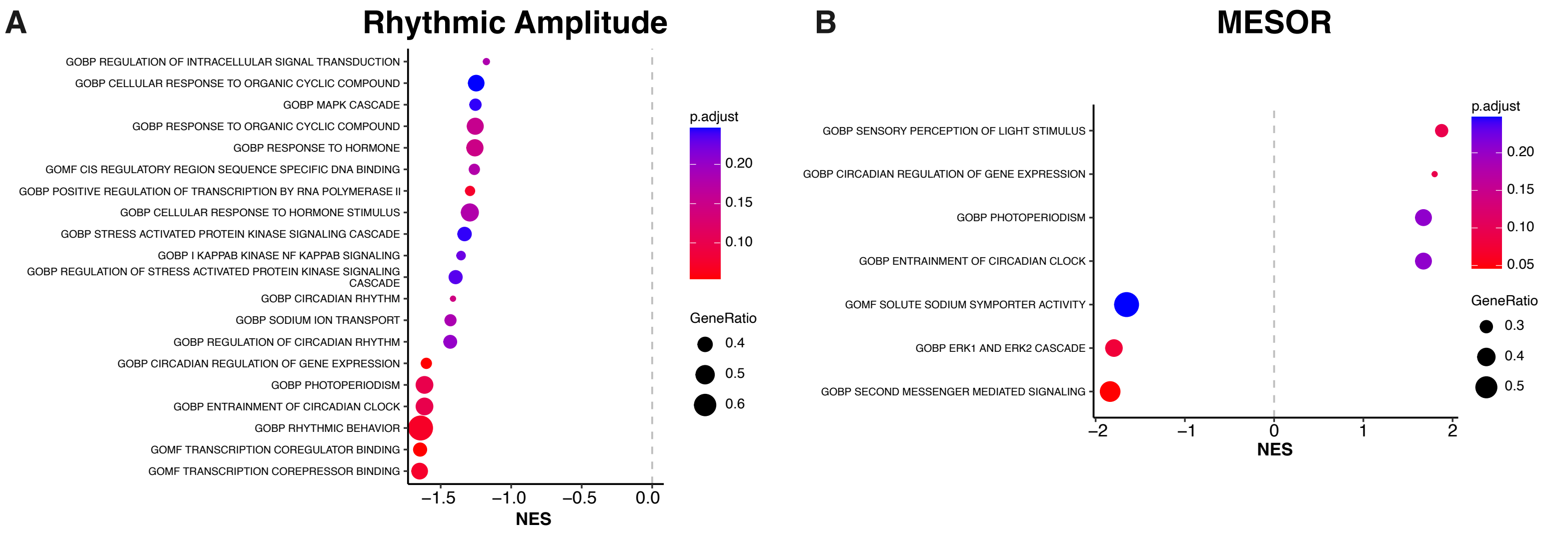


**Supplementary Figure 3. Gene set enrichment analysis (GSEA) for p-value significant rhythmic amplitude and MESOR genes in DiffCircaPipeline . A)** Pathway analysis showing relevant enriched gene sets (p-adj < 0.25) for genes with differential rhythmic amplitude ($p < 0.05$). **B)** Pathway analysis showing relevant enriched gene sets (p-adj < 0.25) for genes with differential MESOR ($p < 0.05$).

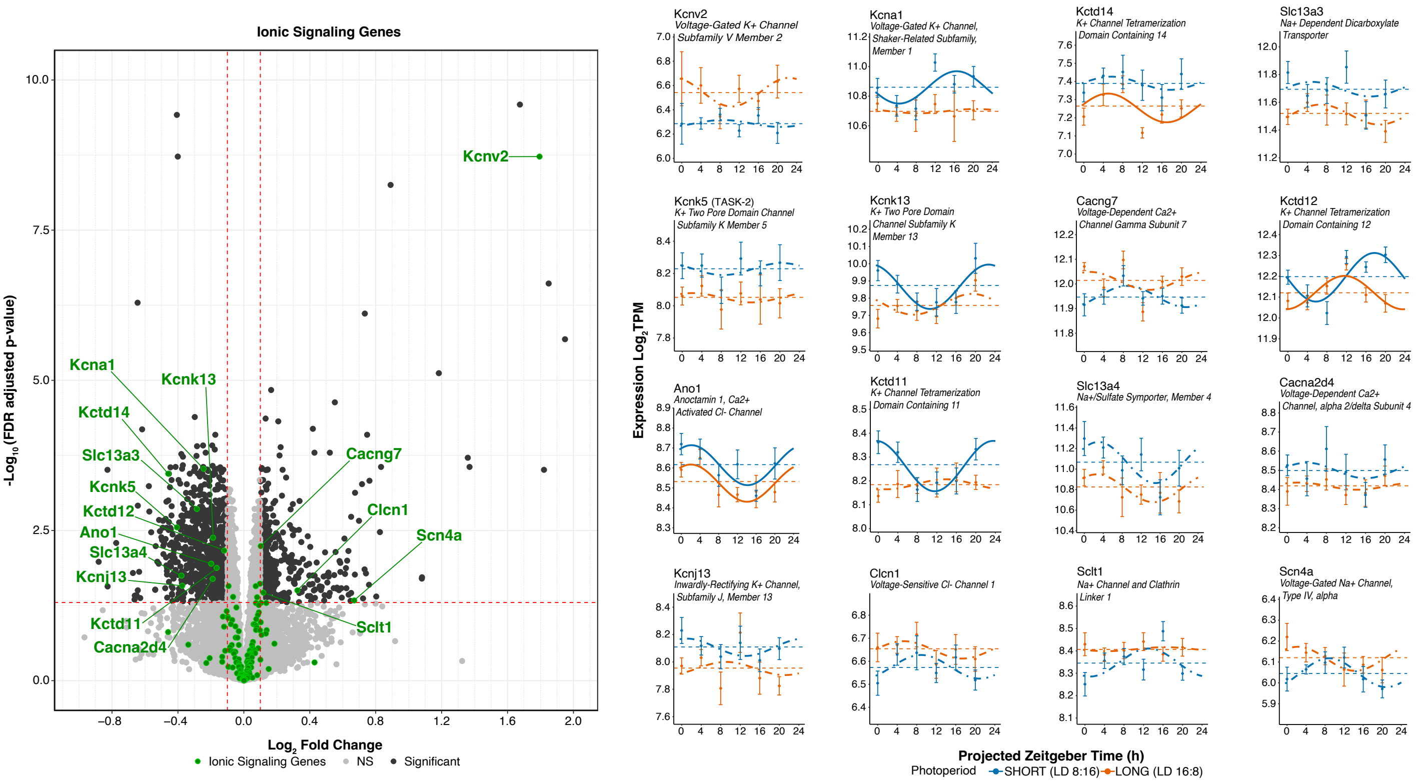


**Supplementary Figure 4. Characterizing ionic signaling gene expression & rhythmicity in response to photoperiod. A)** DESeq2 volcano plot of gene expression highlighting ionic signaling genes in green. Genes meeting the $\log_2 FC \geq \pm 0.1$ and p-adj < 0.05 are labeled. **B)** Cosinor plots of differentially expressed genes. Solid lines depict significantly rhythmic cosinor fits, whereas dot-dashed traces indicate non-rhythmic fits.

Dots represent mean ±SEM for samples collected at each timepoint (n>7). Short (8:16) and Long (16:8) photoperiods are represented in blue and orange, respectively.

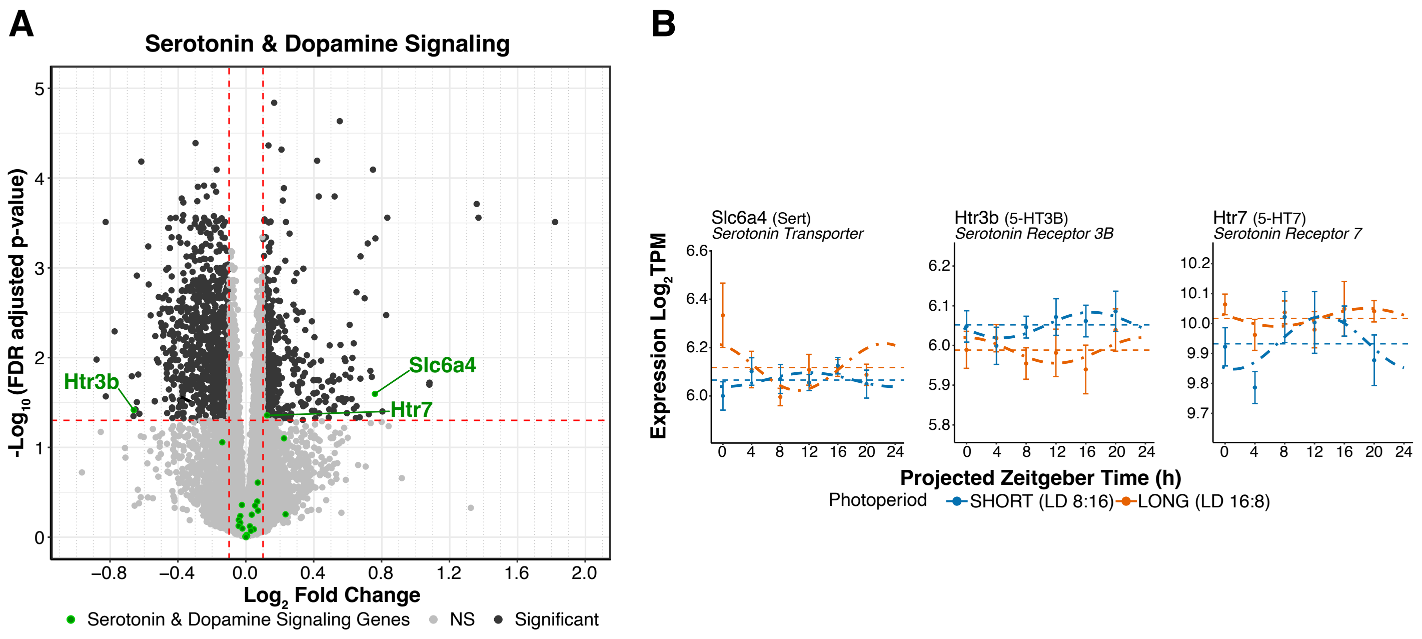


**Supplementary Figure 5. Characterizing monoamine signaling gene expression & rhythmicity in response to photoperiod. A)** DESeq2 volcano plot of gene expression highlighting monoamine signaling genes in green. Genes meeting the $log_2FC \geq \pm0.1$ and p-adj < 0.05 are labeled. **B)** Cosinor plots of differentially expressed genes. Solid lines depict significantly rhythmic cosinor fits, whereas dot-dashed traces indicate non-rhythmic fits.

Dots represent mean ±SEM for samples collected at each timepoint (n>7). Short (8:16) and Long (16:8) photoperiods are represented in blue and orange, respectively.
